# Resveratrol attenuates high-fat diet-induced obesity and the aging-related sarcopenia mitochondrial dysfunction in skeletal muscle

**DOI:** 10.1101/823088

**Authors:** Chyi-Huey Bai, Javad Alizargar, Jia-Ping Wu

**Author notes:** **Correspondence author:** Jia-Ping Wu, Research Center for Healthcare Industry Innovation, National Taipei University of Nursing and Health Sciences. No. 365, Mingde Rd., Beitou Dist., Taipei City 11219, Taiwan, R.O.C.

## Abstract

Sarcopenic obesity is a progressive loss of skeletal muscle mass and strength with increases in adiposity. The aim of this study was to investigate the effects of resveratrol on obesity or sarcopenia to potential therapy risk for skeletal muscle declines in physical function. C57BL/6J male mice were fed either a high-fat diet for 4 weeks and resveratrol (low-, middle-, and high-dose) for 8 weeks to express the obesity effect. Samp8 mice sarcopenia skeletal muscle functional deterioration expressed an age-associated decline. Resveratrol (150 mg/Kg BW, daily) was administered by oral gavage two times a week one month of the experimental period. Exercise training based on adaptations in the muscle is training twice a week for 4 weeks. The skeletal muscles from mice in each group were analyzed by H&E staining, TUNEL and western blot analysis to determine mitochondrial function expression, apoptosis and relative fibrosis signaling. Results of the present study indicate that resveratrol in obesity skeletal muscle is linked to an increase in the expression of mitochondrial function involved in Bcl-2 and PI3K/AKT. On the other hand, resveratrol attenuates sarcopenia Samp8 mice, the age-related loss of skeletal muscle mass and mitochondrial function involved in Bad, caspase 3 and IL-6/ERK1. However, exercise training not find a significant difference in sarcopenia skeletal muscles SAMP8 mice. Exercise training didn’t induce sarcopenia skeletal muscle hypertrophy in sarcopenic SAMP8 mice. Therefore, we suggest that resveratrol as a therapeutic potential in the combination of sarcopenia and obesity, the state called sarcopenic obesity.

## Introduction

Sarcopenic obesity is the combination of sarcopenia and obesity. Some key age- and obesity-mediated factors and pathways may aggravate sarcopenia. Sarcopenia is an age-associated skeletal muscle mass loss coupled with functional deterioration exacerbated by obesity (1). To reflect the direction of the pathological pathway from “sarcopenia→obesity” to “obesity → sarcopenia” be defined as “Sarcopenic obesity”. Over Adipocytes in skeletal muscle tissue are leading to premature tissue aging (2,3). Obesity in adulthood is a medical condition in which excess body fat has accumulated leading to reduced lower skeletal muscle mass and strength with potential implications for developing sarcopenic obesity increased health problems (4,5–6). We hypothesize that resveratrol treatment in male mice programmed by obesity-induced sarcopenia and aging-associated sarcopenia would prevent skeletal muscle atrophy and muscle mass decreases. The prevalence of obesity in the whole word is increasing in all age groups. Since sarcopenia is directly attributed to obesity or age, there’s no guarantee that a person will be able to prevent it (7,10–11). Obesity is most commonly caused by a combination of excessive food energy intake, not exercising enough, age, smoking or heavy alcohol use, although a few caused by genes (12). Obesity is associated with skeletal muscle loss and impairgenesis. Increased infiltration of proinflammatory in skeletal muscle strophy is noted in obesity and sarcopenia and is associated with muscle dysfunction (13,14–16). Sarcopenia is now recognized as a disease, most common in older people, but can also occur earlier in life (17). As we grow older, up to half of the muscle is lost and skeletal muscle is often replaced with fat tissue, particularly in sarcopenia. There are several factors contributing to the disorder such as lack of exercise, and poor nutrition (18). Resveratrol is a phytoalexin polyphenolic compound derived from naturally grapes, red wine and other food products with antioxidant activities and antiaging properties (19,20). Resveratrol rely on into the blood circulation of trans-reveratrol to achieve. Resveratrol has beneficial effects on enhance the complex deficiency of immune system, correction the biochemical defect in oxygen consumption, reduce the risk of cardiovascular disease, inhibition inflammation and platelet aggregation and improve degenerative Alzheimer’s disease (21,22–24). In this study, we investigated the peripheral immune-regulating potential of resveratrol against HFD-induced apoptosis by restoring mitochondrial functional activities, and stimulating mitochondrial biogenesis. Resveratrol and exercise training is a highly effective strategy to offset sarcopenia (25). Exercise training improves the vasodilatory properties of the vasculature thereby optimizing O_2_ transport throughout the body to improve vascular function in association with the reduction of reactive oxygen species (26). Blood flow is markedly increased in contracting skeletal muscles and myocardium. Habitual exercise training alone did not elicit mitochondrial dysfunction improvements in healthy aged subject, may be have to combined with resveratrol intake in aging-associated function decline (27, 28–32). Therefore, we determine the effect of resveratrol, exercise training, and combined exercise training with resveratrol intake on age-dependent sarcopenia. And to determine the effect of resveratrol on obesity-induced sarcopenia or age-dependent sarcopenia in sarcopenic obesity health.

## Materials and methods

### Obesity animal model

We purchased 46 male C57BL/6J mice at ages 6 weeks from BioLASCO Taiwan Co., Ltd. C57BL/6J mice (N=39) were fed a high-fat diet (HFD) for 5 weeks to induce obesity, and then were assigned to three groups containing low- (50 mg/Kg BW, N=10), middle-(100 mg/Kg BW, N=10) and high- (200 mg/Kg BW, N=9) dose of resveratrol and 0.2 % pubucol (N=10) for 4 weeks. The HFD group received a high-fat diet, and the HFD+LR group received a low-dose resveratrol (LR)-containing HFD, the HFD+MR group received a middle-dose resveratrol (MR)-containing HFD, the HFD+HR group received a high-dose resveratrol (HR)-containing HFD and 0.2 % pubucol-containing HFD. After additional feeding of the experimental diet for 4 weeks, mice in the HFD group were highly obese compared with the mice in the standard diet fed mice group. The use of animals in our experimental design was approved by the Animal Care and Use Committee of Taipei Medical University. Adult mice were housed five per cage in a temperature-controlled room (24 ± 1℃) with a 12-h light/dark cycle (06.30a.m. to 18.30p.m.). Mice were fed commercial laboratory chow and water ad. lib. Animals were killing after 4 week experimental. The skeletal muscles were collected for dissection of muscle mass and detected the skeletal muscle to quantify mitochondrial function and aging-related disease signaling proteins by Western blotting. We also dissected and weighed of white adipose tissue to analyze adiposity, besides body protein body fat (%) content analysis by muscle drying weight (mg)/fat weight (mg). All experimental procedures were approved by the Animal Care and Use Committee of Taipei Medical University, and all animal experiments were performed in accordance with the ARRIVE guidelines and use of laboratory animals.

### Experimental design in sarcopenia animal model

Threety-one male adult senescence-accelerated mice (SAMP8) of 3 months of age weighting 35 g were used in this study (total N=31). We examined the effect of exercise training, resveratrol and their combination on the prevention of sarcopenia using a senescence-accelerated-prone mice (SAMP8) model. Animals were divided into four experimental groups: non-treated SAMP8 control groups (SAMP8, N=8), exercise training SAMP8 groups (SAMP8+Ex, N=8), resveratrol intake SAMP8 groups (SAMP8+ Re, 150 mg/Kg BW, N=10) (N=7) and combination exercise training and resveratrol intake SAMP8 groups (SAMP8+Ex+Re, N=8). Mice were housed in two cage of one group for all experimental process before sacrificed. Mice maintained in a room at 22±2 °C, with automatic light cycles (12-h light/dark) and standard diet ad libitum. All the animals received animal care according to the Guidelines for Ethical Care of the Animal Care and Use Committee of Taipei Medical University (LAC-2019-0264). Mice were exercise training involved in protein synthesis and degradation were upregulated in the skeletal muscles, suggesting accelerated protein turnover. Total body and adipose tissue weights decreased following the use of the exercise training cage. Thus, physical exercise training may assist in recovering from aging-related sarcopenia (loss of muscle function) and obesity in SAMP8 mice. All animal studies are complied with the ARRIVE guidelines.

### Hematoxylin-eosin stained (H&E) and terminal deoxynucleotidyl transferase dUTP nick end labeling (TUNEL) positive nuclei stain

After skeletal muscle perfusion fixation with buffered formalin for 15min, the skeletal muscle was sectioned into six equal transverse slices starting from the apex to base. Two of these sections were embedded in paraffin. The remaining sections were cut at 6μm thickness and stained with H&E stain. Slides were deparaffinized and dehydrated. Samples were passed through a series of graded alcohols from 100% to 90% to 70%, 5min each. The skeletal muscle sections were processed for fibrosis quantification. H&E staining were prepared one of two sections and incubated for 5min at room temperature. The other of two sections were treated to TUNEL assay (Apoptosis detection kit, TA300, R&D Systems, Minneapolis, MN, USA). After rinsing with phosphate-buffered saline (PBS), each slide was then soaked with 85% alcohol, 100% alcohol for 15min. Stained sections were rinsed with PBS and air dried before mounting. Cross-sections were H&E stained and TUNEL assay. Color images of cross-sections were made at 200x total magnification using a Nikon E600. Microscope lighting was optimized and increased the probability of being visualized in appropriate cross-sections and not-tangentially. The number of TUNEL-positive cells per muscle area was counted in 20 visual fields (magnification 400×) for each mice.

### Western blotting analysis

The skeletal muscle cut in small pieces in PBS buffer homogenized in a homogenizer and then centrifuged at 12,000 ×rpm for 30min at 4°C. Supernatants were considered total cellular cytosol lysates protein. Separate the protein samples using gel electrophoresis. SDS-PAGE was carried out with 10~12.5% polyacrylamide gels. The protein were electrophoresed at 100V for 1.5hr. Electrophoresis proteins were transferred to PVDF membranes (PVDF membrane, 0.45μm; MILLIPORE, USA) using a Bio-Red Laboratories Instruments Mini Trans-Blot Electrophoretic Transfer Cell unit (Alfred Nobel Drive Hercules, California, USA) at 150mA for 2hr in transfer buffer (25mM Tris-HCl, 192mM glycine and 20% methanol. pH8.3). We blocked PVDF membranes in 5% non-fat milk buffer (diluted in Tris-buffered saline and 0.1% Tween 20) for 1hr at room temperature and then in blocking buffer containing 100mM Tris-HCl, pH 7.5, 0.9% NaCl, and 0.1% fetal bovine serum for 2hr at room temperature. For transfer buffers without methanol it is essential that complete equilibration of the resolving gel is achieved to prevent distortions within the gel which would cause band smearing. Only a brief rinse is required to achieve equilibration if the transfer buffer contains methanol. Monoclonal antibodies (Santa Cruz Biotechnology, Santa Cruz, CA, USA) were diluted 1:250 in antibody binding buffer containing 100mM Tris-HCl, pH 7.5, 0.9% NaCl, 0.1% Tween-20, and 1% fetal bovine serum. Incubations were performed at 4 °C for 2hr. Western blot analysis was performed as previously described (Sibilia et al., 2000) with antibodies detecting ANP, BNP, FGF-2, UPA, PI3K, AKT, IL-6, STAT3, p-JNK, p-P38, and tubulin (Santa Cruz). Bcl-2, Bad, Cytochrome c, caspase 3, MMP2, and MMP9 (Cell Signaling). NFACc3 and ERK1 (Abcam). Washed the immunoblots three times in 5ml blotting buffer for 5min and afterwards PVDF were incubated with secondary antibody solution containing Horseradish peroxidase- conjugated anti-rabbit (1:1000), anti-goat (1:500), or anti-mouse (1:1000) IgG secondary antibodies (Santa Cruz Lab, CA) for 1hr at room temperature. Secondary antibodies were used for ECL chemiluminescent (Sigma-Aldrich, Bornem, Belgium) detection of proteins. When required, blots were reprobed after stripping in 62.5mM Tris-HCl (pH 6.8), 2%SDS, and 100 mMβ-mercaptoethanol at room temperature for 30min, blocked and reprobed with polyclonal or monoclonal antibody. Membranes were scanned and quantified with Image J Imagine program software.

### Statistical analysis

Quantitation was carried out by analyzing the intensity of the hybridization signals using ImageJ Imagine program for western blot analysis. Obesity (HFD-fed mice) data was obtained from at least two independent experiments. Statistical analysis of the data was normalized to the values of the group of the same batch expressed as a mean ± standard error of the means (SEM) using SigmaStat software. \**p* < 0.05, \*\**p* < 0.01 significant differences compared with control group, ^#^*p* < 0.05, ^##^*p* < 0.01 significant differences compared with SAMP8 without treatment group. Sarcopenia (SAMP8 mice) data was obtained from at least three independent experiments. Statistical analysis of the data was performed to the values of the control group of the same batch using SigmaStat 11.0 software. Results were expressed as mean ± SEM. Two-group comparisons were carried out with the Student’s t test. Differences were considered significant at \**p* < 0.05 and \*\**p* < 0.01.

## Results

### Resveratrol prevents age-related changes in the body and skeletal muscle composition in sarcopenia in adults with obesity and SAMP8 mice

In adults with obesity, sarcopenia is associated with lower muscle mass and strength and a greater likelihood of occurrence sarcopenia obesity. C57BL/6J mice fed high-fed diet (HFD) to induce obesity, body drying weight (mg), muscle drying grinding (mg), fat weight (mg) and body fat (%) were investigated. Results showed body drying weight-HFD groups mean ± SEM. (6.98417 ± 1.135715 (mg), p=0.030959) was increased compared to control (5.94437 ± 0.70812 (mg)). Resveratrol intake during HFD-fed was no significant difference compared to HFD groups. Compared to control, all of dose resveratrol lead to body drying weight increases (Table 1). Bodyweight drying weight is skeletal muscle and fat couple together. Skeletal muscle drying grinding-HFD groups mean ± SEM. (6.98417 ± 1.135715 (mg), p=0.040394) was increased compared to control (5.76293 ± 0.74955 (mg)). High-dose resveratrol containing HFD (HFD+HR (200 mg/Kg)) groups mean ± SEM. (7.01058 ± 0.59543 (mg), p=0.14457) was a little skeletal muscle increases compared to HFD groups (6.59688 ± 0.912178 (mg)) (Table 2). In contrast, low-dose resveratrol containing HFD (HFD+LR (50 mg/Kg)) (1.30442 ± 0.420315 (mg), p=0.058043) groups and middle-dose resveratrol containing HFD (HFD+MR (100 mg/Kg)) (1.11554 ± 0.451432 (mg), p=0.017555) of fat weight (mg) has significant difference decreases, when compared to HFD (1.76943 ± 0.73303 (mg)) groups (Table 3). HFD+LR (%) (19.9915 ± 5.560475 (mg), p=0.046902) groups and HFD+MR (17.40719 ± 5.992501 (%), p=0.013303) of body fat (%) has significant difference decreases, when compared to HFD (%) (1.76943 ± 0.73303 (%) groups (Table 4). Taken a together, in adulthood obesity independent predictor of lower skeletal muscle mass and high-fat weight with potential implication for sarcopenia in adults with obesity. Resveratrol can make skeletal muscle mass increases and lower fat weight during sarcopenia in adults with obesity development. On aging-related skeletal muscle atrophy, we use senescence-accelerated mice (SAMP8) to investigate the growing recognition of sarcopenia. Results showed in SAMP8 mice, the age-related loss of skeletal muscle mass, exercise training, resveratrol intake, and their combination grow or maintain skeletal muscle mass (Fig. 1A). Age-related sarcopenia induced skeletal muscle fiber disorder in SAMP8 mice. Exercise training, resveratrol intake, and their combination ameliorated the increase in slow skeletal muscle fibers in the SAMP8 mice (Fig. 1B and 1C). Sarcopenia recognized muscle-wasting condition is suffering from a major of cell apoptosis. Cell apoptosis was observed using TUNEL staining (Fig. 1D). Exercise training, resveratrol intake, and their combination alleviated cell apoptosis in skeletal muscle sarcopenic SAMP8 mice (Fig. 1D and 1E). We demonstrate for the first time that resveratrol has a therapeutic effect against skeletal muscle sarcopenia.

**Table 1.** Body drying weight (mg) of the skeletal muscle in HFD-fed C57BL/6J mice. HFD: High-fat diet; LR: Low–dose resveratrol; MR: Middle– dose resveratrol; HR: High–dose resveratrol. Statistical analysis of the data was performed the values using SigmaStat 11.0 software. Results are expressed mean ± SE. Groups comparison to control or HFD were carried out with the Student’s t test. Differences were expressed at *p* value.

| Body drying weight (mg) | control | HFD |  |  |  |
| --- | --- | --- | --- | --- | --- |
|  |  | HFD | LR | MR | HR |
| N | 7 | 10 | 10 | 10 | 9 |
| mean | 5.94437 | 6.98417 | 6.60851 | 6.43085 | 7.27751 |
| SEM | 0.70812 | 1.135715 | 0.590994 | 0.608673 | 0.71902 |
| p. value |  |  |  |  |  |
| v.s. control |  | 0.030959 | 0.033646 | 0.087547 | 0.0019 |
| v.s. HFD |  |  | 0.19516 | 0.106987 | 0.26907 |

**Table 2.** Skeletal muscle dring grinding (mg) of the skeletal muscle in HFD-fed C57BL/6J mice. HFD: High-fat diet; LR: Low–dose resveratrol; MR: Middle–dose resveratrol; HR: High–dose resveratrol. Statistical analysis of the data was performed the values using SigmaStat 11.0 software. Results are expressed mean ± SEM. Groups comparison to control or HFD were carried out with the Student’s t test. Differences were expressed at *p* value.

| Skeletal muscle drying grinding (mg) | control | HFD | LR | HFD<br>MR | HR |
| --- | --- | --- | --- | --- | --- |
| N | 7 | 10 | 10 | 10 | 9 |
| mean | 5.76293 | 6.59688 | 6.43525 | 6.2767 | 7.01058 |
| SEM. | 0.74955 | 0.912178 | 0.53798 | 0.516849 | 0.59543 |
| p. value |  |  |  |  |  |
| v.s. control |  | 0.040394 | 0.03068 | 0.068493 | 0.00187 |
| v.s. HFD |  |  | 0.32626 | 0.185844 | 0.14457 |

**Table 3.**
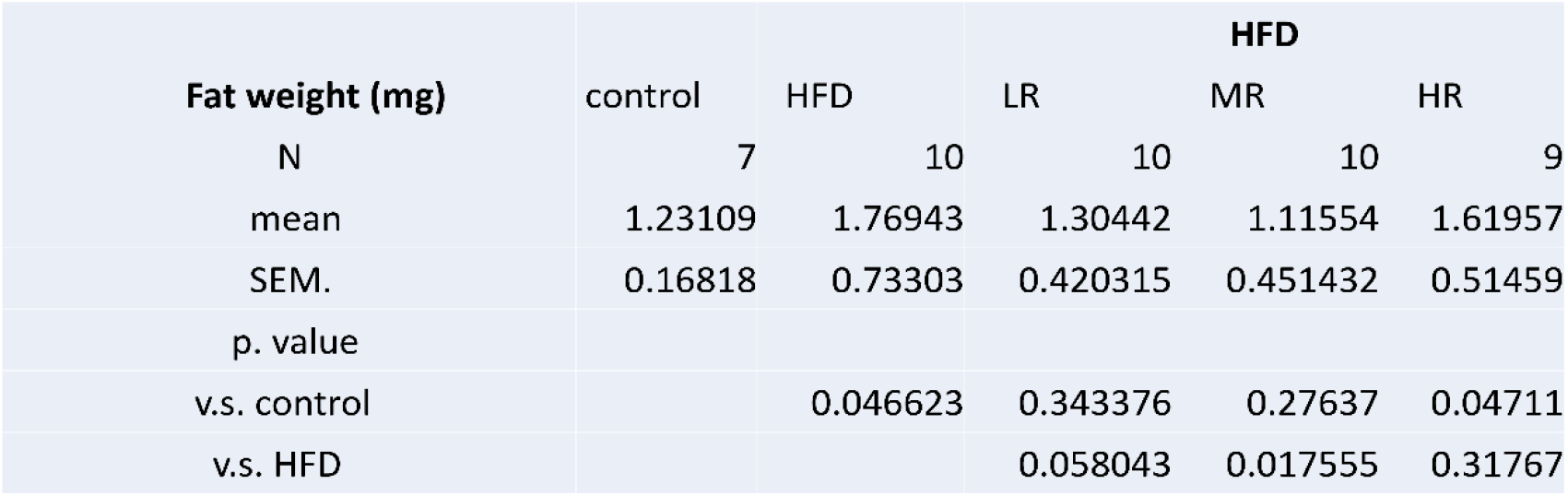
Fat weight (mg) of the skeletal muscle in HFD-fed C57BL/6J mice. HFD: High-fat diet; LR: Low–dose resveratrol; MR: Middle–dose resveratrol; HR: High–dose resveratrol. Statistical analysis of the data was performed the values using SigmaStat 11.0 software. Results are expressed mean ± SEM. Groups comparison to control or HFD were carried out with the Student’s t test. Differences were expressed at *p* value.

**Table 4.** Body fat (%) of the skeletal muscle in HFD-fed C57BL/6J mice. HFD: High-fat diet; LR: Low–dose resveratrol; MR: Middle–dose resveratrol; HR: High–dose resveratrol. Statistical analysis of the data was performed the values using SigmaStat 11.0 software. Results are expressed mean ± SEM. Groups comparison to control or HFD were carried out with the Student’s t test. Differences were expressed at *p* value.

|  |  | HFD |  |  |  |
| --- | --- | --- | --- | --- | --- |
| **Body fat (%)** | control | HFD | LR | MR | HR |
| N | 7 | 10 | 10 | 10 | 9 |
| mean | 21.42796 | 26.58601 | 19.9915 | 17.40719 | 22.76621 |
| SEM. | 1.757741 | 9.7018368 | 5.560475 | 5.992501 | 5.63407 |
| p. value |  |  |  |  |  |
| v.s. control |  | 0.1055508 | 0.272062 | 0.063088 | 0.289916 |
| v.s. HFD |  |  | 0.046902 | 0.013303 | 0.170754 |

**Figure 1.**
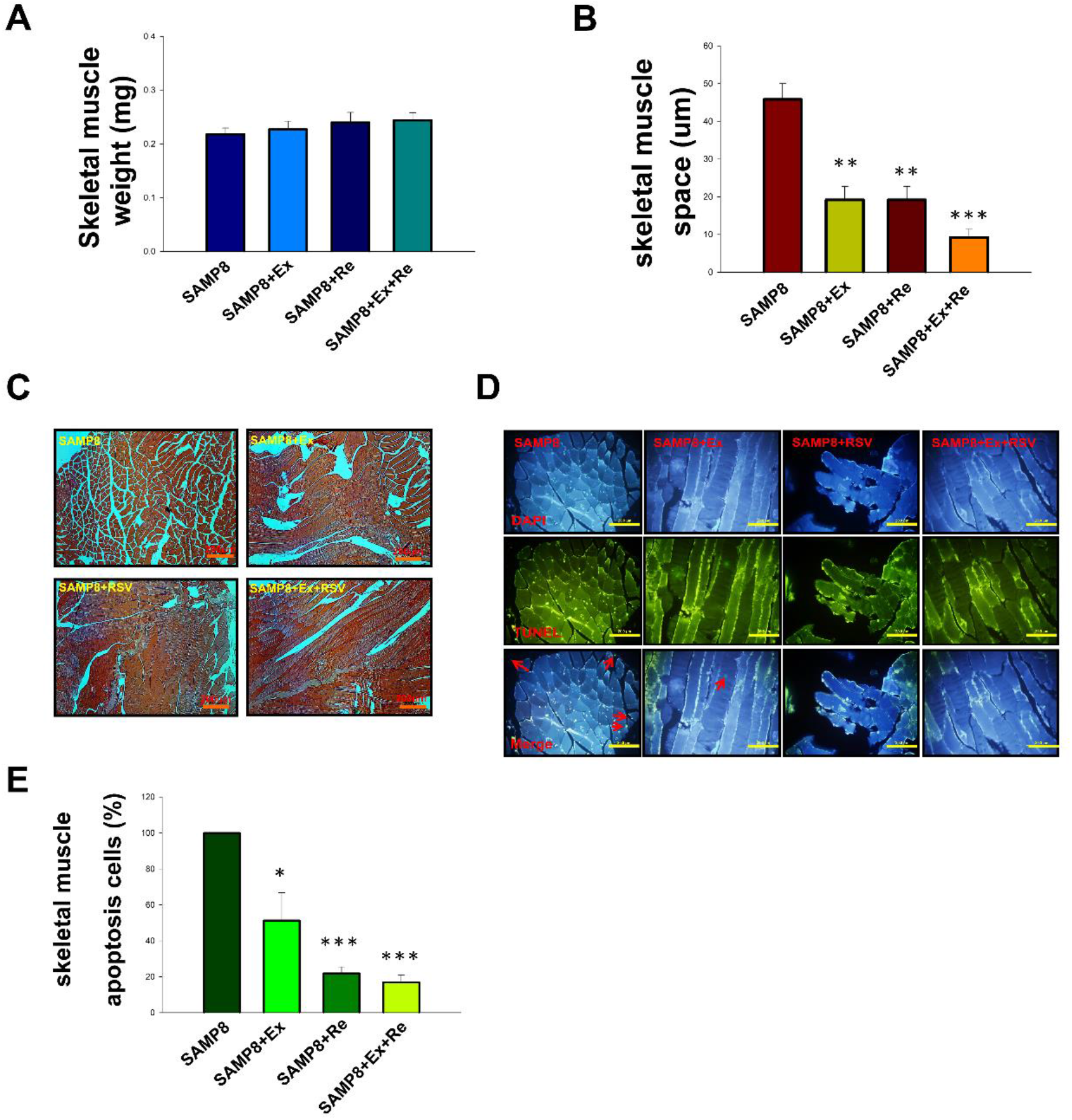
Sarcopenia is the age-related loss of muscle mass and function in SAMP8 mice. (A). Skeletal muscle weight (mg) in exercise training, resveratrol intake combination of exercise training and resveratrol intake in SAMP8 mice. (B). Skeletal muscle space (um) in exercise training, resveratrol intake combination of exercise training and resveratrol intake in SAMP8 mice. (C). Histology of Skeletal muscle in exercise training, resveratrol intake combination of exercise training and resveratrol intake in SAMP8 mice using H&E staining. (D). Apoptosis cells of Skeletal muscle in exercise training, resveratrol intake combination of exercise training and resveratrol intake in SAMP8 mice using TUNEL staining. (E). Statistical analysis results is expressed skeletal muscle apoptosis cells (%). All data is expressed mean ± SEM. ^*^*p*<0.05, ^**^*p*<0.01, \*\*\**p*<0.001, significant difference compared with SAMP8 mice without treatment. SAMP8; SAMP8 control groups, SAMP8+Ex; exercise training SAMP8 groups, SAMP8+ Re; resveratrol intake SAMP8 groups, SAMP8+Ex+Re; combination exercise training and resveratrol intake SAMP8 groups.

### Resveratrol counteracts obesity and aging-associated decline of mitochondrial biogenesis and function in skeletal muscles of HFD-fed mice and sarcopenic skeletal muscles SAMP8 mice

Consumption of high-fat diet (HFD) is related with increased dysfunctional mitochondria in skeletal muscles. Thus, in order to determine mitochondria function, we detected mitochondrial protein populations in high-fat diet (HFD)-induced obesity mice. Results showed us high-fat diet (HFD)-induced obesity augments skeletal muscle atrophy by induction of protein degradation in the mitochondria (Fig. 2). High-dose of the anti-adipogenic agents resveratrol preserved mitochondrial activity in skeletal muscle from mice with high-fat diet (HFD)-induced obesity (Fig. 2A). High-dose of resveratrol has a significant difference compared to control or HFD-fed mice by upregulation of Bcl 2 and downregulation of Bad protein expression (vs. control, p<0.05 and vs. HFD, p<0.05). Pubucol also be found the same results in Bal 2 and Bad. These findings suggest that the HFD-induced loss of mitochondria and HFD-diet induced impairment of mitochondrial function may combine to promote skeletal muscle atrophy. Age-related skeletal muscle weakness, sarcopenia, was detected mitochondrial function proteins, Bcl 2, Bad, Cytochrome and Caspase 3, in sarcopenic skeletal muscle SAMP8 mice (Fig. 2B). No significant change was observed for Bcl 2, Bad, Cytochrome c, and Caspase 3 protein expression levels of exercise training in sarcopenic skeletal muscle SAMP8 mice. Results supporting therapeutic effects of resveratrol in the sarcopenia muscle atrophy, we found Bcl 2 increases and Bad, Cytochrome c, and Caspase 3 decreases. Especially, combination exercise training and resveratrol intake has a significant increases by Bcl 2 (p<0.01), in contrast, decreases by Bad (p<0.01), Cytochrome c (p<0.01), and Caspase 3 (p<0.05) in sarcopenic skeletal muscle SAMP8 mice.

**Figure 2.**
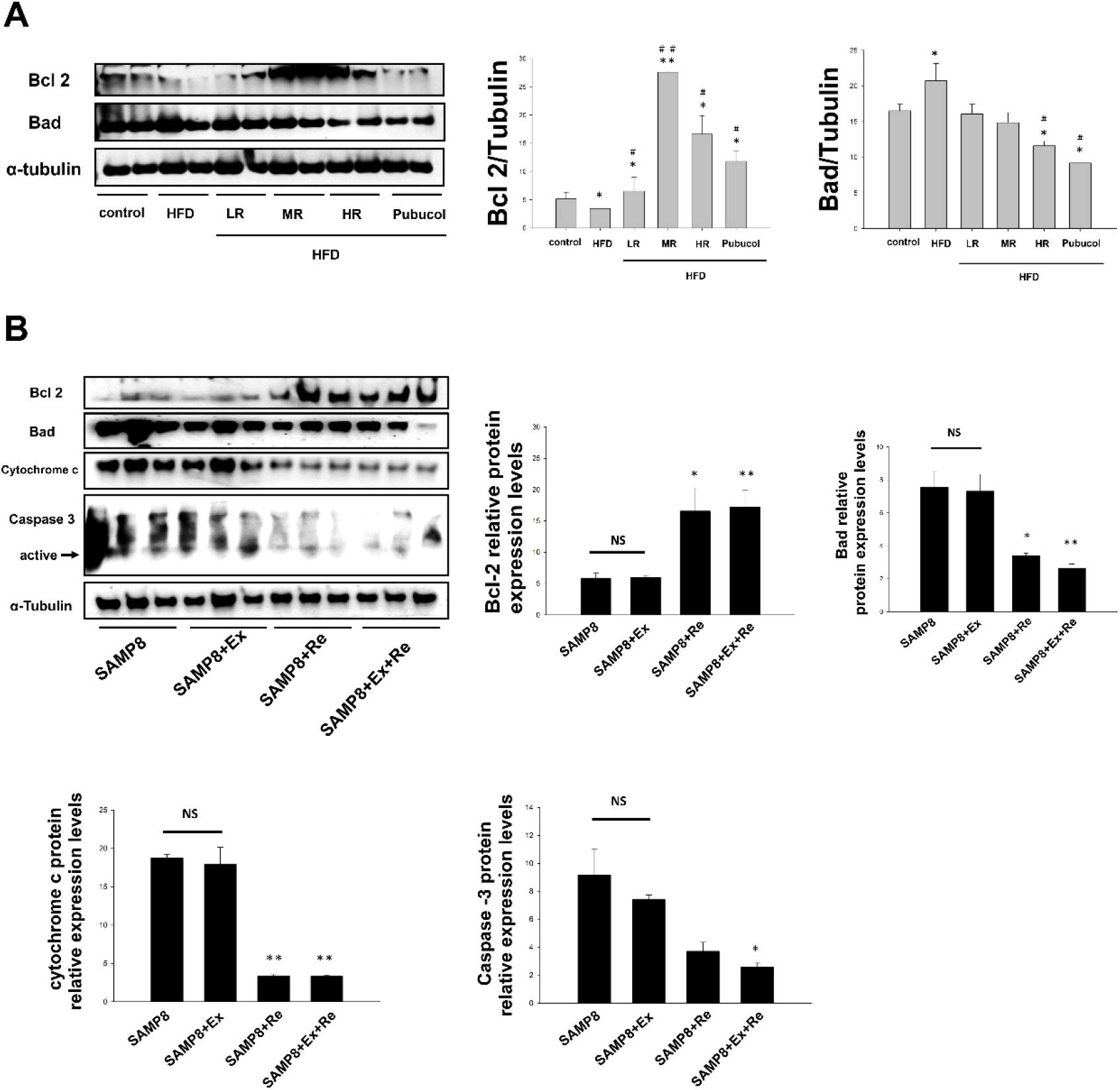
Resveratrol modulates the mitochondrial dysfunction to apoptosis in skeletal muscles of obesity-induced sarcopenia and age-related sarcopenia SAMP8 mice. (A). Representative western blotting analysis for Bcl 2 and Bad protein levels by high-fat diet (HFD)-induced obesity. All data is expressed mean ± SEM. \**p*<0.05, ^**^*p*<0.01, significant difference compared with control. ^#^*p*<0.05, ^##^*p*<0.01, significant difference compared with HFD mice without treatment. α-tubulin was used as a loading control. HFD: High-fat diet; LR: Low–dose resveratrol; MR: Middle–dose resveratrol; HR: High–dose resveratrol. (B). Representative western blots using antibodies against Bcl 2, Bad, Cytochome c, Caspase 3, and α-tubulin in sarcopenia SAMP8 mice. After quantification, Bcl 2/α-tubulin, Bad/α-tubulin, Cytochrome c/α-tubulin, and Caspase 3/α-tubulin ratios were calculated in sarcopenia SAMP8 mice. α-tubulin was used as a loading control. Data are expressed as mean ± SEM. ^*^*p*<0.05, ^**^*p*<0.01, significant difference compared with SAMP8 mice control. NS. was no significant difference. SAMP8; SAMP8 control groups, SAMP8+Ex; exercise training SAMP8 groups, SAMP8+ Re; resveratrol intake SAMP8 groups, SAMP8+Ex+Re; combination exercise training and resveratrol intake SAMP8 groups

### Resveratrol promoted protein synthesis during the muscle hypertrophic process following atrophy reveals dramatic increases in sarcopenia extracellular matrix remodeling

MMP2 and MMP9 play a role in adipose and muscle hypertrophy and could be involved in ECM (extracellular matrix remodeling) in response to HFD diet. Skeletal muscle ECM remodeling occurs to HFD-induced obesity with as skeletal muscle mass gain (Table 1 and Table 2). MMP2 and MMP9 up-regulation enhanced activity in obesity skeletal muscle atrophy (p<0.05). Resveratrol is only slightly elevated in MMP9 HDF-induced skeletal muscle (Fig. 3A). Low-dose resveratrol-containing HFD (HFD+LR) and middle-dose resveratrol-containing HFD (HFD+MR) groups has a significant difference compared to control (p<0.05). MMP9 up-regulation appears to be an important additional molecular event in the multistep process of all inflammatory myopathies. The results provide a potential interventional strategy for controlling inflammatory response by supplementing resveratrol. In adipose tissue, atrial natriuretic peptide (ANP) and brain natriuretic peptide (BNP) increase the release of free fatty acids from adipose tissue. Obesity in adipose tissue induced by high-fat diet is associated with impaired natriuretic peptide release leading to a significant reduction of systemic natriuretic peptide expression levels. In skeletal muscle tissue, high-fat diet is a little difference increases by ANP and BNP. The actions of BNP are mediated via the ANP receptors, the physiologic effects of BNP are identical to those of ANP. Resveratrol intake lower ANP and BNP protein synthase was found to be a predictor of survival in high-fat diet-induced obesity sarcopenia skeletal muscle (Fig. 3B). Resveratrol was minimally altered by HFD-diet induced obesity sarcopenia. MMP2 and MMP9 are involved in aging physiological processes such as skeletal muscle development and maturation. Degeneration of skeletal muscle MMP2 and MMP9 with age contributed to functional decline in sarcopenia SAMP8 mice (Fig. 3C). Exercise training did not reverse these changes. In order for exercise training to be effective, proper nutrition must be in place. Resveratrol intake plays a major role in preventing sarcopenia. Results showed resveratrol intake and exercise training supplements resveratrol intake can contributed skeletal muscle MMP2 and MMP9 with age increases (p<0.05 and p<0.01, respectively). Furthermore, to show the potential for fibroblast growth factor (FGF2) and Urokinase-type plasminogen activator (UPA) to affect skeletal muscles. Exercise training remained FGF2 and UPA activities unchanged. FGF2 is involved in muscle mass modulation. Both growth factors of FGF2 and UPA activities enhance the reinnervation of muscle. Thus, exercise training, resveratrol intake, and exercise training supplements resveratrol intake no effects on FGF2 and UPA activities (Fig. 3D). Exercise training and resveratrol intake treatment with FGF2 and UPA causes skeletal muscle hypertrophy in SAMP8 mice.

**Figure 3.**
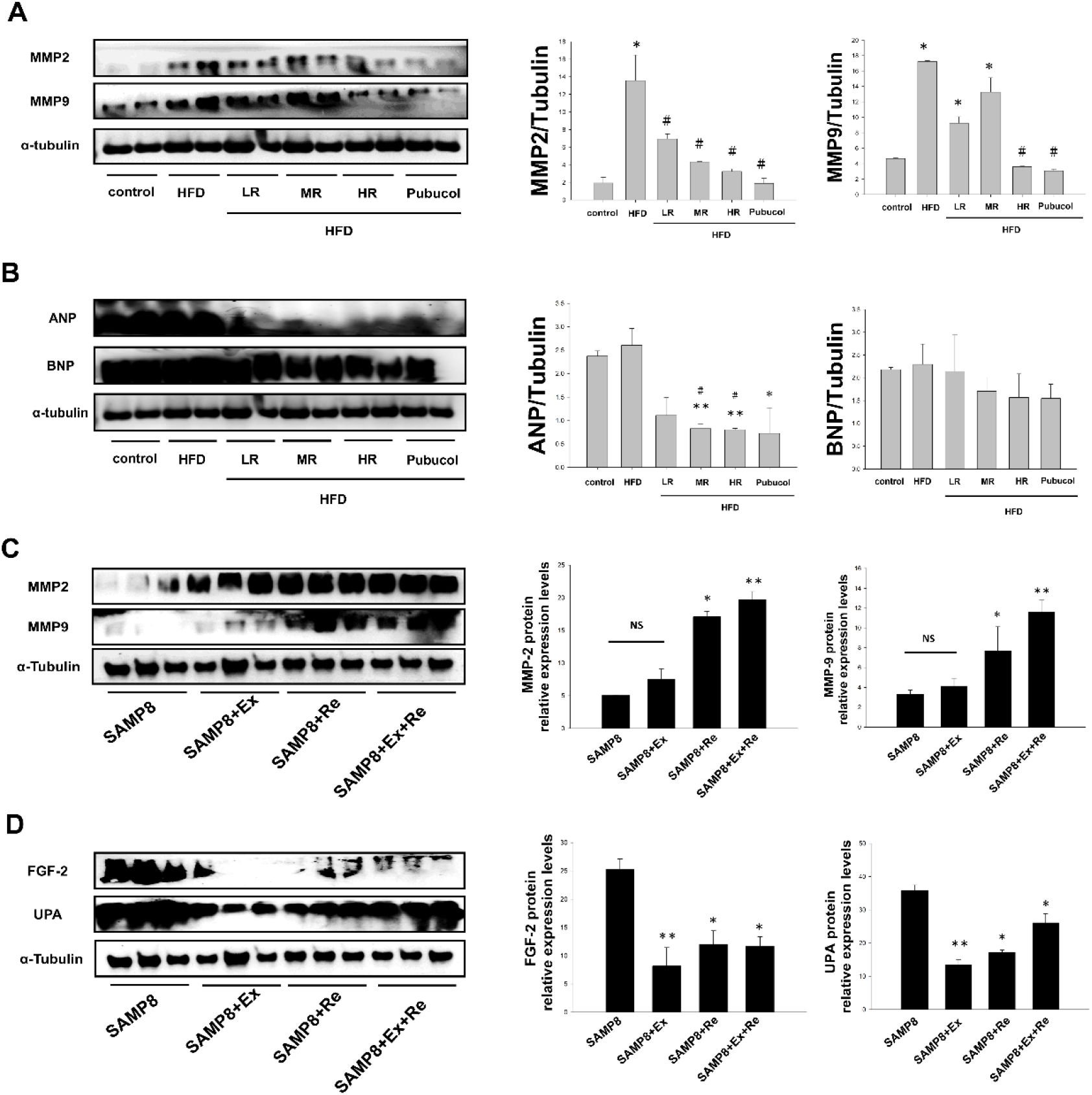
Resveratrol reduces matrix metalloproteinases in skeletal muscles of obesity-induced sarcopenia, but elevates matrix metalloproteinases in age-related sarcopenia SAMP8 mice. (A). The levels of MMP2, MMP9, ANP, and BNP protein in high-fat diet (HFD)-induced obesity were measured by western blotting. All data is expressed mean ± SEM. ^*^*p*<0.05, ^**^*p*<0.01, significant difference compared with control. ^#^*p*<0.05, significant difference compared with HFD mice without treatment. α-tubulin was used as a loading control. HFD: High-fat diet; LR: Low–dose resveratrol; MR: Middle–dose resveratrol; HR: High–dose resveratrol. (B). Western blot analysis of MMP2, MMP9, FGF-2, and UPA were performed in sarcopenia SAMP8 mice. α-tubulin was used as a loading control. Quantification of Western blot bands by densitometry. All data are expressed as mean ± SEM. ^*^*p*<0.05, ^**^*p*<0.01, significant difference compared with SAMP8 mice without treatment. NS. was no significant difference. SAMP8; SAMP8 control groups, SAMP8+Ex; exercise training SAMP8 groups, SAMP8+ Re; resveratrol intake SAMP8 groups, SAMP8+Ex+Re; combination exercise training and resveratrol intake SAMP8 groups.

### Resveratrol protected against skeletal muscle atrophy and inflammatory myopathies induced by obesity and sarcopenia

High-fat diet (HFD) induced obesity has deleterious effect on skeletal muscle atrophy and is associated with low-grade chronic inflammation. On the molecular level, HFD-induced obesity inhibited PI3K/Akt pathway in skeletal muscles of HFD-fed mice (p<0.05). Resveratrol intake in spite of low-, middle-, or high-dose was activated PI3K/Akt pathway in skeletal muscles of HFD-fed mice (Fig 4A). Resveratrol intake was to be a promising material to alleviate HFD-induced muscle atrophy. Obesity is associated with chronic inflammation after adjustment for aging. Chronic inflammation occurs in skeletal muscle in obesity is mainly manifested by increased immune cell infiltration and proinflammatory activation in intermyocellular and perimuscular adipose tissue. Thus, skeletal muscle inflammation showed distinct patterns of up-regulation of IL-6 and STAT3 in skeletal muscles of HFD-fed mice (Fig. 4B) (p<0.05). High-dose resveratrol in HFD-fed mice (HR-HFD) and Pubucol has significant lower expression levels by IL-6 (p<0.05). STAT3 was a slight increase in skeletal muscle inflammation. Thus, resveratrol in HFD-fed mice (HR-HFD) has no significant difference by STAT3. Taken together, resveratrol alleviated adulthood obesity-induced sarcopenia in skeletal muscles of HFD-fed mice. Furthermore, we determine whether resveratrol could alleviate age-related sarcopenia in skeletal muscle SAMP8 mice. Age-associated with sarcopenia is a disease of muscles in which skeletal muscle atrophy and inflammatory myopathies. Age-associated with sarcopenia can be slightly mitigated, if not outright managed, with exercise and nutrition. Intriguingly, young adult SAMP8 mice with slight sarcopenia were no effects by exercise training. In this study, we only find resveratrol has good effects against skeletal muscle atrophy and inflammatory myopathies (Fig. 4C and 4D). We found protein expression of PI3K and Akt increases by resveratrol (p<0.05) and combination of exercise training and resveratrol (PI3K, p<0.05 and Akt, p<0.01) in sarcopenic skeletal muscles SAMP8 mice (Fig. 4C). Interestingly, down regulation of IL-6 and STAT3 was found by resveratrol and combination of exercise training and resveratrol. IL-6 protein expression was decreased by resveratrol (p<0.05) and combination of exercise training and resveratrol (p<0.01). STAT3 was slight decreases by combination of exercise training and resveratrol in sarcopenic skeletal muscles SAMP8 mice (Fig. 4D). Obesity and inflammation are reportedly associated with pathogenesis of sarcopenia. Mice showed increased skeletal muscle p38 MAPK and JNK activities and were resistant to the development of HFD fed-induced obesity by resveratrol in spite of dose (Fig. 5A). Skeletal muscle pro-inflammation promote obesity, in part, by activating the p38 MAPK (p<0.05) and JNK. In pro-inflammation skeletal muscle of obesity, JNK and p38 MAPK is upregulated in HFD-fed mice which resolved their paradoxical activation during diseased and healthy conditions. Resveratrol increased skeletal muscle p38 MAPK and JNK activities and were resistant to the development of diet-induced obesity. Thus, activation of the p38 MAPK/JNK module promotes obesity. In young SAMP8 mice despite unchanged the p38 MAPK/JNK protein expression stimulated by exercise training and their combination (Fig. 5B). Resveratrol stimulated p38 MAPK promotes prevent skeletal muscle atrophy (p<0.05). Thus, low-grade systemic inflammation is a component of skeletal muscle pathologies (sarcopenia) can be prevented by performing regularly physical exercise. Sarcopenia caused by aging factor, but the development of it can be attributed to skeletal muscle mass loss resulting in skeletal mass atrophy. Therefore, we detected skeletal muscle hypertrophy factors, MFACc3 and ERK1/2, in skeletal muscles of HFD-fed mice and sarcopenic skeletal muscles SAMP8 mice. We were investigated ERK1/2 and NFATc3 in HFD-fed induced obesity mice and sarcopenia SAMP8 mice of skeletal muscle hypertrophy or atrophy. Results has shown that ERK1/2 and NFATc3 protein expression was decreases in HFD-fed induced obesity mice and sarcopenia SAMP8 mice of skeletal muscle hypertrophy (p<0.05).

**Figure 4.**
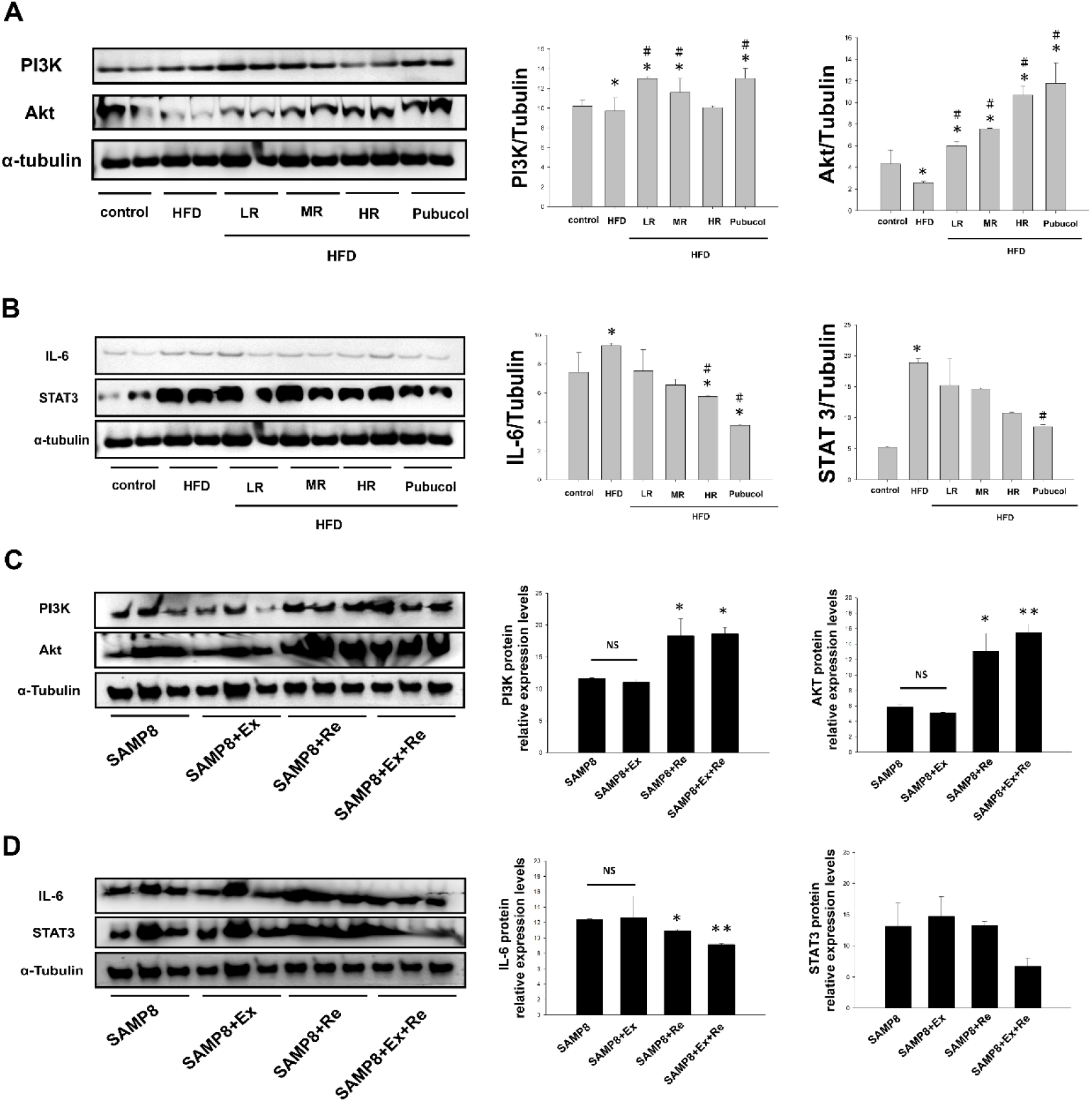
Effects of resveratrol on the recovery of skeletal muscle anti-inflammatory and anti-atrophy properties in skeletal muscles of obesity-induced sarcopenia and age-related sarcopenia SAMP8 mice. (A). Western blotting for anti-inflammatory (IL/STAT3) and anti-atrophy (PI3K/Akt) properties in high-fat diet (HFD)-induced obesity. α-tubulin was used as a loading control. Quantitative analysis of PI3K, Akt, IL-6, and STAT3 levels. All data is expressed mean ± SEM. ^*^*p*<0.05, significant difference compared with control. ^#^*p*<0.05, significant difference compared with HFD mice without treatment. α-tubulin was used as a loading control. HFD: High-fat diet; LR: Low–dose resveratrol; MR: Middle– dose resveratrol; HR: High–dose resveratrol. (B). Western blotting of PI3K, Akt, IL-6, and STAT3 protein levels in sarcopenia SAMP8 mice. α-tubulin was used as a loading control. Quantitative analysis of PI3K, Akt, IL-6, and STAT3 levels. Data are expressed as mean ± SEM. ^*^*p*<0.05, ^**^*p*<0.01, significant difference compared with SAMP8 mice without treatment. NS. was expressed no significant difference. SAMP8; SAMP8 control groups, SAMP8+Ex; exercise training SAMP8 groups, SAMP8+ Re; resveratrol intake SAMP8 groups, SAMP8+Ex+Re; combination exercise training and resveratrol intake SAMP8 groups.

**Figure 5.**
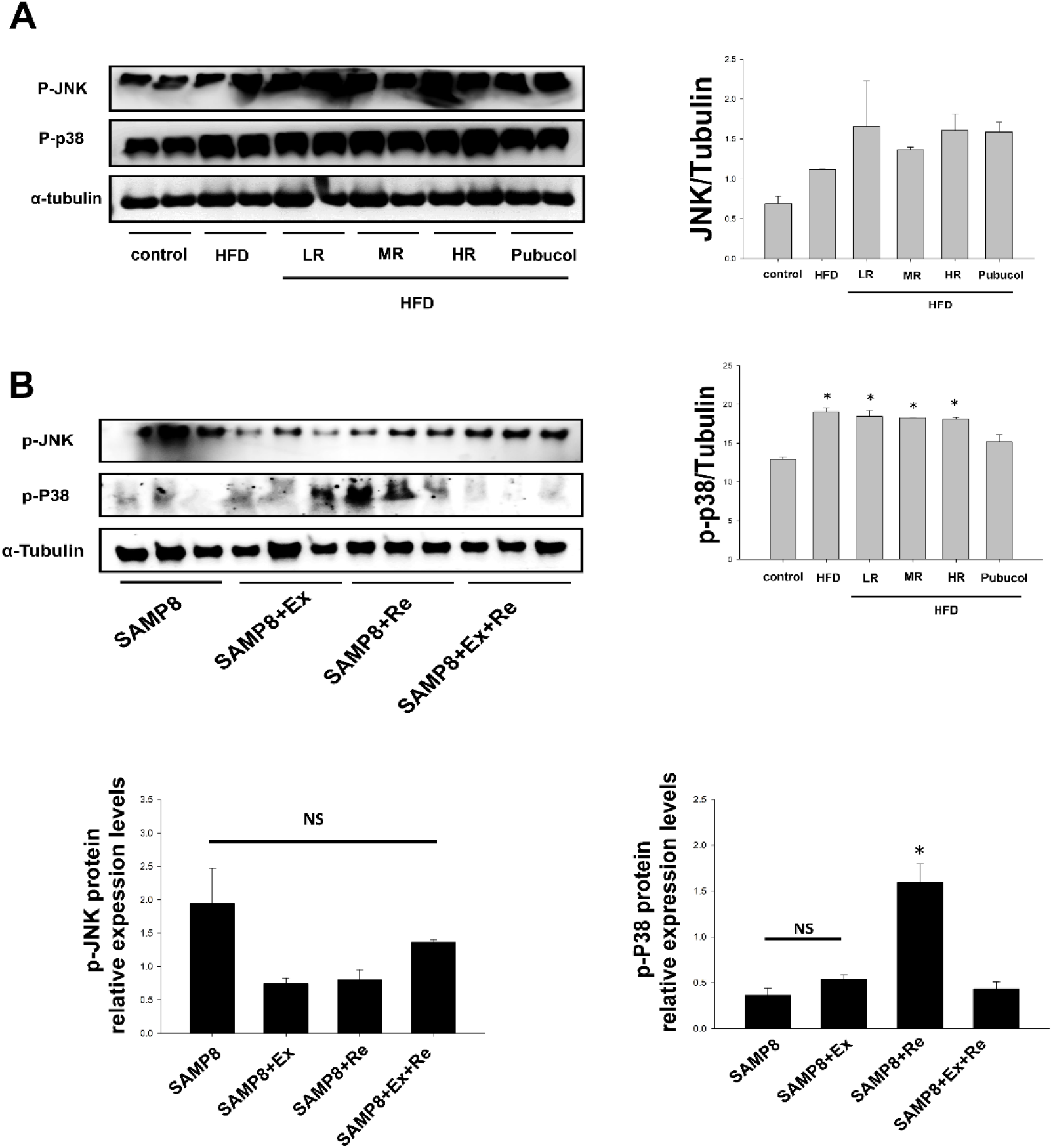
Resveratrol induces skeletal muscle p38 MAPK and JNK activities to resistant the skeletal muscles of obesity-induced sarcopenia and reduced JNK in aging-related sarcopenia SAMP8 mice. (A).Western Blotting analysis of skeletal muscle pro-inflammation showed increased levels of p-JNK and p-38 in high-fat diet (HFD)-induced obesity. α-tubulin was used as a loading control. All data is expressed mean ± SEM. ^*^*p*<0.05, significant difference compared with control. α-tubulin was used as a loading control. HFD: High-fat diet; LR: Low– dose resveratrol; MR: Middle–dose resveratrol; HR: High–dose resveratrol. (B).Representative western blotting analysis of p-JNK and p-P38 proteins in skeletal muscle sarcopenia SAMP8 mice after exercise training, resveratrol intake or their combination are shown. α-tubulin was used as a loading control. Data are expressed as mean ± SEM. ^*^*p*<0.05, significant difference compared with SAMP8 mice without treatment. NS. was no significant difference. SAMP8; SAMP8 control groups, SAMP8+Ex; exercise training SAMP8 groups, SAMP8+ Re; resveratrol intake SAMP8 groups, SAMP8+Ex+Re; combination exercise training and resveratrol intake SAMP8 groups.

Resveratrol prevented the exacerbation of skeletal muscle atrophy (Fig. 6A). After intake in spite of low-, middle-, and high- dose resveratrol, NFATc3 was a little increase to cause hypertrophy and prevent atrophy. In contrast, ERK1/2 was not activated during HFD-induced skeletal muscle atrophy (Fig. 6B) (p<0.05). Low-, middle-, and high- dose resveratrol, activation of ERK1/2 can oppose skeletal muscle atrophy induced by disuse. For skeletal muscle facing aging populations, sarcopenia lead to a loss in the mass and size of our muscles in SAMP8 mice. Sarcopenia has been associated with impaired skeletal muscle hypertrophy. Exercise training a little reverse muscle atrophy occurred to NFATc3 increases (Fig. 6C). Resveratrol and their combination was not found good effects on skeletal muscle hypertrophy or atrophy. In sarcopenic skeletal muscle SAMP8 mice, ERK1 protein expression did not have significant difference by exercise training (Fig. 6D). Resveratrol and their combination has significant difference by ERK1 (p<0.05). Exercise and proper nutrition help battle this disease. Therefore, resveratrol protected against skeletal muscle atrophy and inflammatory myopathies induced by obesity and sarcopenia.

**Figure 6.**
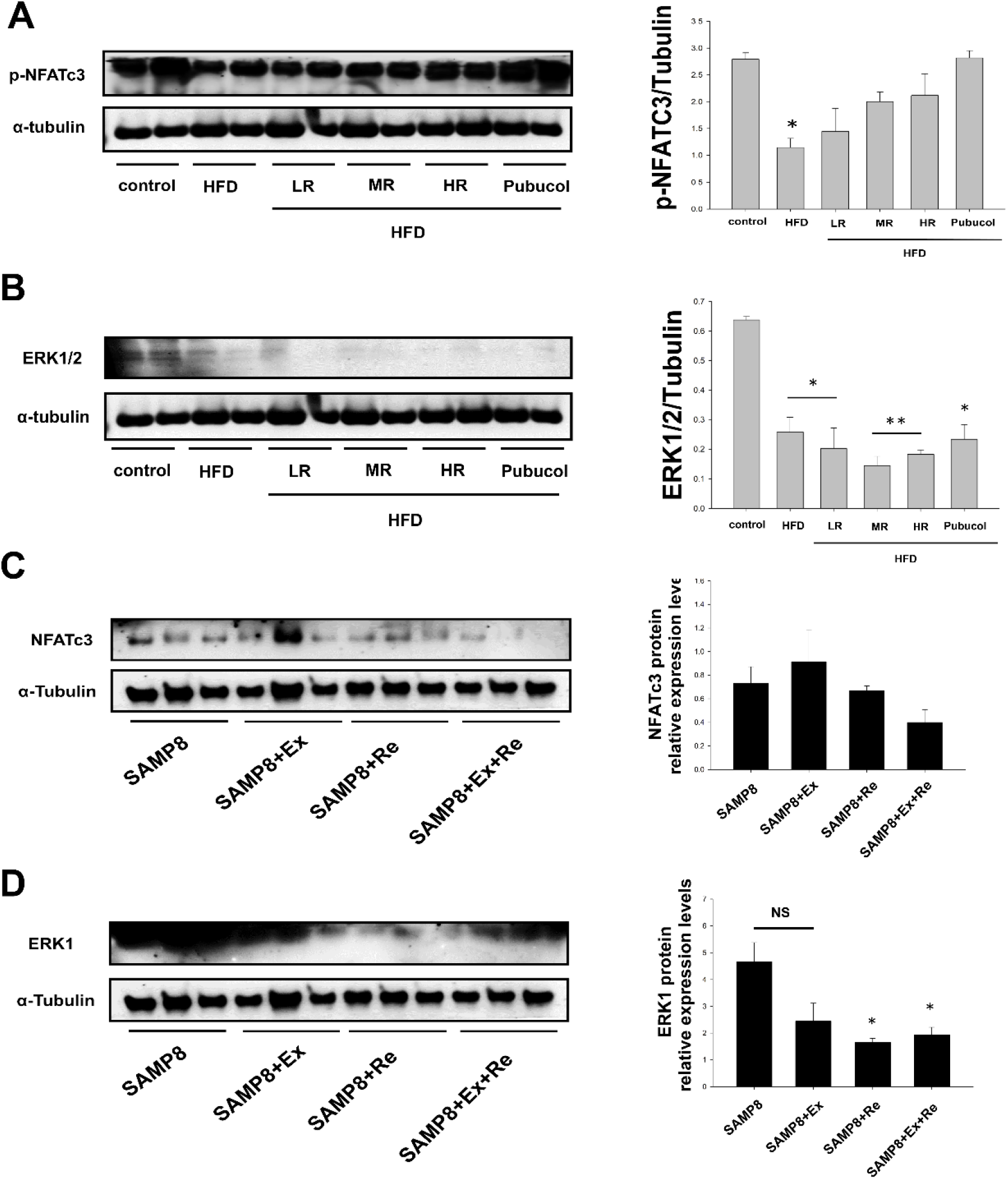
Resveratrol causes skeletal muscle hypertrophy in protecting muscle from atrophy in skeletal muscles of obesity-induced sarcopenia and age-related sarcopenia SAMP8 mice. (A). Western blotting of NFATc3 and ERK1/2 protein levels in high-fat diet (HFD)-induced obesity. All data is expressed mean ± SEM. \**p*<0.05, ^**^*p*<0.01, significant difference compared with control. α-tubulin was used as a loading control. HFD: High-fat diet; LR: Low–dose resveratrol; MR: Middle–dose resveratrol; HR: High–dose resveratrol. (B). Protein levels of NFATc3 and ERK1 by Western blotting in sarcopenia SAMP8 mice. α-tubulin was used as a loading control. Data are expressed as mean ± SEM. ^*^*p*<0.05, significant difference compared with SAMP8 mice without treatment. NS. was no significant difference. SAMP8; SAMP8 control groups, SAMP8+Ex; exercise training SAMP8 groups, SAMP8+ Re; resveratrol intake SAMP8 groups, SAMP8+Ex+Re; combination exercise training and resveratrol intake SAMP8 groups.

## Discussion

The aging of the population represents one of the largest healthcare challenges facing the world today. Some papers has reported to lower the rate of production of free radical reaction damage have been shown to increase the life span, to enhance mitochondrial function and age-mediated immune responses and to slow development of sarcopenia and the skeletal muscles disorders (33, 34–37). We examined the effects of resveratrol, a natural polyphenolic compound present in grapes, on sarcopenia obesity in (high-fat diet (HFD)-induced obesity mice and aging-associated sarcopenia SAMP8 mice. The resveratrol theory of aging provides reasonable explanations for age-associated sarcopenica (38). In addition, studies strongly suggest that resveratrol play a significant role in the deterioration of skeletal muscle with obesity and sarcopenia with age-associated phenomena (39). It was found in this study that HFD-fed mice with resveratrol during obesity-sarcopenia could improve skeletal muscle mass (Table 1 and Table 2) and inhibited fat weight (Table 3 and Table 4). Adipose tissue extra lipids cannot be accumulated in the adipose tissue much longer, does infiltrate into peripheral organs such as the skeletal muscle cause dysfunction of the organs (40,41). Deleterious effects of aging probably reflecting on skeletal muscle mass (Fig. 1A), space (Fig. 1B and 1C), and cell apoptosis (Fig. 1D and 1E). Furthermore, expression levels of mitochondrial biogenesis and function, muscle hypertrophic process following atrophy, and skeletal muscle atrophy and inflammatory myopathies in skeletal muscle tissue from two different mice models of high-fat diet (HFD)-induced obesity mice and aging-associated sarcopenia SAMP8 mice were detected by western blot. Resveratrol has been reported to target mitochondrial-related pathways in two different animal models (Fig. 2 and Fig. 3). Exercise training does not play a role in regulating skeletal muscle mitochondrial dysfunction, inflammatory myopathies, and muscle hypertrophic process during aging-related atrophy (sarcopenia) (Fig. 2, Fig. 3, Fig. 4, Fig. 5 and Fig. 6). In general, resveratrol has also been shown to exhibit antioxidant, anti-inflammatory, anti-proliferative properties and to decrease collage synthesis in two skeletal muscle type animal models. Therefore, resveratrol is currently advised as supplement in the diet of elderly individuals. Sarcopenic obesity results in skeletal muscle and fat replacement, resveratrol which prevent skeletal muscles atrophy from obesity or sarcopenia (42). Resveratrol induces mediates anti-inflammatory effects and inhibits skeletal muscles atrophy. (Fig. 4 and Fig. 5). Moreover, matrix metalloproteinases (MMPs) in inflammatory myopathies enhanced immunoreactivity near atrophic myofibers (43). MMP-9 was strongly expressed in atrophic myofibers in all inflammatory myopathies. MMP-2 immunoreactivity is only slightly elevated in inflamed muscle. (Fig. 3A and 3C). ANP and BNP binding to the natriuretic peptide receptor A (NPRA) rising intracellular cGMP levels induce expression of downstream genes in skeletal muscle (44). Free fatty acids from adipocyte lipolysis serve as additional ligands for the transcriptional regulator of mitochondrial biogenesis. Thus, resveratrol increase skeletal muscle mitochondrial content (Bcl2, PI3K/Akt) (Fig. 2 and Fig. 4) and skeletal muscle reverse atrophy, but not hypertrophy (FGF2/UPA and NFATc3/ERK1/2) (Fig. 3 and Fig. 6). A novel function of FGF-2 in regulating skeletal muscle mass through enlargement of muscle fiber size, while physiological and pharmacological doses, treatment with FGF-2 causes skeletal muscle hypertrophy in mice, to protect muscle from atrophy (45, 46–47). JNK/p38-mediated intrinsic pathway signaling is one of the mechanisms involved in age-related increase in muscle cell apoptosis (48, 49-50). Protective effect of resveratrol against inflammation on progressive skeletal muscle atrophy also be observed in obesity-induce sarcopenia mice and age-related sarcopenia SAMP8 mice.

## Abbreviation

MMP 9: Matrix metalloproteinases 9
MMP 2: Matrix metalloproteinases 2
HFD: High-fat dairy
ANP: Atrial natriuretic peptide
BNP: Brain natriuretic peptide
PI3K: phosphatidylinositol 3 kinase
JNK: c-Jun NH2-terminal kinase
NFATC3: Nuclear factor of activated T-cells
uPA: Urokinase-type plasminogen activator
SAMP8: senescence-accelerated prone mice

## Author contributions

Conceptualization: J.P. Wu., C.H. Bai, J. Alizargar; Methodology: C.H. Bai, J. Alizargar, J.P. Wu; Formal analysis: C.H. Bai, J. Alizargar, J.P. Wu; Investigation: C.H. Bai, J. Alizargar, J.P. Wu; Resources: C.H. Bai, J. Alizargar, J.P. Wu; Data curation: J.P. Wu, J. Alizargar; Writing original draft: J.P. Wu; Writing review and editing: J.P. Wu; Visualization: J.P. Wu; Supervision: C.H. Bai, J. Alizargar, J.P. Wu; Project administration: J.P. Wu; Funding acquisition: J.P. Wu

## Acknowledgements

Not applicable.

## References

1. Haslam, D. (2017). Men and obesity: what are the issues? Trends in Urology & Men’s Health. 8, 21–24.

2. Haslam, D. (2008). Understanding obesity in the older person: prevalence and risk factors. Br. J. Community. Nurs. 13, 115–11 probably reflecting denervated 6.

3. Denison, H. J., Cooper, C., Sayer, A. A. and Robinson, S. M. (2015). Prevention and optimal management of sarcopenia: a review of combined exercise and nutrition interventions to improve muscle outcomes in older people. Clin. Interv. Aging. 10, 859–869.

4. Laurentius, T., Kob, R., Fellner, C., Nourbakhsh, M., Bertsch, T., Sieber, C. C. and Bollheimer, L. C. (2019). Long-Chain Fatty Acids and Inflammatory Markers Coaccumulate in the Skeletal Muscle of Sarcopenic Old Rats. Disease Markers. 1–11.

5. Heo, J. W., Yoo, S. Z., No, M. H., Park, D. H., Kang, J. H., Kim, T. W. Kim, C. J. Seo, D. Y., Han, J. and Yoon, J. H. et al. (2018). Exercise Training Attenuates Obesity-Induced Skeletal Muscle Remodeling and Mitochondria-Mediated Apoptosis in the Skeletal Muscle. Int J Environ Res. Public. Health.15, 2301.

6. Chen, H. T., Chung, Y. C., Chen, Y. J., Ho, S. Y. and Wu, H. J. (2017). Effects of Different Types of Exercise on Body Composition, Muscle Strength, and IGF-1 in the Elderly with Sarcopenic Obesity. Journal of the American Geriatrics Society. 65, 827–832.

7. Aubrey, J., Esfandiari, N., Baracos, V. E., Buteau, F. A., Frenette, J., Putman, C. T. and Mazurak, V. C. (2014). Measurement of skeletal muscle radiation attenuation and basis of its biological variation. Acta Physiologica. 210, 489–497.

8. Lettieri-Barbato, D., Cannata, S. M., Casagrande, V., Ciriolo, M. R. and Aquilano, K. (2018). Time-controlled fasting prevents aging-like mitochondrial changes induced by persistent dietary fat overload in skeletal muscle. PLoS One. 13, e0195912.

9. Choi, W. H., Son, H. J., Jang, Y. J. Ahn, J., Jung, C. H. and Ha, T. Y. (2017) Apigenin Ameliorates the Obesity-Induced Skeletal Muscle Atrophy by Attenuating Mitochondrial Dysfunction in the Muscle of Obese Mice. Mol Nutr Food Res. 61.

10. Frias, F. T., Rocha, K. C. E., de Mendonça, M., Murata, G. M., Araujo, H.N., de Sousa, L. G. O., de Sousa, É., Hirabara, S. M., Leite, N. C. and Carneiro, E. M., et al. (2018). Fenofibrate reverses changes induced by high-fat diet on metabolism in mice muscle and visceral adipocytes. J Cell Physiol. 233, 3515–3528.

11. Chen, G., Ye, G., Zhang, X., Liu, X., Tu, Y., Ye, Z. Liu, J., Guo, Q., Wang, Z. and Wang, L., et al. (2018). Metabolomics Reveals Protection of Resveratrol in Diet-Induced Metabolic Risk Factors in Abdominal Muscle. Cellular Physiology & Biochemistry. 45, 1136–1148.

12. Haohao, Z., Guijun, Q., Juan, Z., Wen, K. and Lulu, C. (2015). Resveratrol improves high-fat diet induced insulin resistance by rebalancing subsarcolemmal mitochondrial oxidation and antioxidantion. Physiol Biochem. 71, 121–131.

13. Chen, G., Ye, G., Zhang, X., Liu, X., Tu, Y., Ye, Z., Liu, J., Guo, Q., Wang, Z. and Wang, L., et al. (2018). Metabolomics Reveals Protection of Resveratrol in Diet-Induced Metabolic Risk Factors in Abdominal Muscle. Cell Physiol Biochem. 45, 1136–1148.

14. Kalinkovich, A. and Livshits, G. (2017). Sarcopenic obesity or obese sarcopenia: A cross talk between age-associated adipose tissue and skeletal muscle inflammation as a main mechanism of the pathogenesis. Ageing Res Rev. 35, 200–221.

15. Theodorakopoulos, C., Jones, J., Bannerman, E. and Greig, C. (2017). Effectiveness of nutritional and exercise interventions to improve body composition and musclestrength or function in sarcopenicobese older adults: A systematic review. Nutrition Research. 43, 3–15.

16. Balachandran, A., Krawczyk, S. N., Potiaumpai, M. and Signorile, J. F. (2014). High-speed circuit training vs hypertrophy training to improve physical function in sarcopenic obese adults: a randomized controlled trial. Exp Gerontol. 60, 64–71.

17. Kalinkovich, A., and Livshits, G. (2017). Sarcopenic obesity or obese sarcopenia: A cross talk between age-associated adipose tissue and skeletal muscle inflammation as a main mechanism of the pathogenesis. Ageing Res Rev. 35, 200–221.

18. Kong, D., Song, G., Wang, C., Ma, H., Ren, L., Nie, Q., Zhang, X., and Gan, K. (2013). Overexpression of mitofusin 2 improves translocation of glucose transporter 4 in skeletal muscle of high-fat diet-fed rats through AMP-activated protein kinase signaling. Mol Med Rep. 8, 205–210.

19. Rolland, Y., Lauwers-Cances, V., Cristini, C., van Kan, G. A., Janssen, I. Morley, J. E., and Vellas, B. (2009). Difficulties with physical function associated with obesity, sarcopenia, and sarcopenic-obesity in community-dwelling elderly women: the EPIDOS (EPIDemiologie de l’OSteoporose) Study. AM J CLIN NUTR. 89, 895–1900.

20. Ginés, C., Cuesta, S., Kireev, R., García, C., Rancan, L. Paredes, S. D., Vara, E. and Tresguerres, J. A. F. (2017). Protective effect of resveratrol against inflammation, oxidative stress and apoptosis in pancreas of aged SAMP8 mice. Exp Gerontol. 90, 61–70.

21. Liao, Z.Y., Chen, J. L., Xiao, M. H., Sun, Y., Zhao, Y. X., Pu, D., Lv, A. K., Wang, M. L., Zhou, J. and Zhu, S.Y. et al. (2017). The effect of exercise, resveratrol or their combination on Sarcopenia in aged rats via regulation of AMPK/Sirt1 pathway. Exp Gerontol. 98, 177–183.

22. Kilic, E. M., Kilincli, A. and Eren, Ö. (2015). Resveratrol Induced Premature Senescence Is Associated with DNA Damage Mediated SIRT1 and SIRT2 Down-Regulation. PLoS One. 10, e0124837.

23. Pozo-Guisado, E., Lorenzo-Benayas, M. J., and Fernández-Salguero, P. M. (2004). Resveratrol modulates the phosphoinositide 3-kinase pathway through an estrogen receptor alpha-dependent mechanism: relevance in cell proliferation. Int J Cancer. 109, 167–173.

24. Olesen, J., Gliemann, L., Biensø, R., Schmidt, J., Hellsten, Y., and Pilegaard, H. (2014). Exercise training, but not resveratrol, improves metabolic and inflammatory status in skeletal muscle of aged men. J Physiol. 592,1873–1886.

25. Garcia, P., Schmiedlin-Ren, P., Mathias, J. S., Tang, H., Christman, G. M. and Zimmermann, E. M. (2012). Resveratrol causes cell cycle arrest, decreased collagen synthesis, and apoptosis in rat intestinal smooth muscle cells. Am J Physiol Gastrointest Liver Physiol. 302, G326–35.

26. Franco, J., Dias-Rocha, C., Fernandes, T., Albuquerque Maia, L., Lisboa, P., Moura, E., Pazos-Moura, C. and Trevenzoli, I. (2016). Resveratrol treatment rescues hyperleptinemia and improves hypothalamic leptin signaling programmed by maternal high-fat diet in rats. EUR J NUTR. 55, 601–610.

27. Olesen, J., Ringholm, S., Nielsen, M. M., Brandt, C. T. Pedersen, J. T. Halling, J. F. Goodyear, L. J. and Pilegaard, H. (2013). Role of PGC-1α in exercise training- and resveratrol-induced prevention of age-associated inflammation. Exp Gerontol. 48, 1274–1284.

28. Muhammad, M. H. and Allam, M. M. (2018). Resveratrol and/or exercise training counteract aging-associated decline of physical endurance in aged mice; targeting mitochondrial biogenesis and function. J Physiol Sci. 68, 681–688.

29. Lagouge, M., Argmann, C., Gerhart-Hines, Z., Meziane, H., Lerin, C., Daussin, F., Messadeq, N., Milne, J., Lambert, P. and Elliott, P. et al. (2006). Resveratrol improves mitochondrial function and protects against metabolic disease by activating SIRT1 and PGC-1alpha. Cell. 127, 1109–1122.

30. Fröjdö, S., Durand, C., and Pirola, L. (2008).Metabolic effects of resveratrol in mammals--a link between improved insulin action and aging. Curr Aging Sci. 1, 145–151.

31. Fröjdö, S., Durand, C., Molin, L., Carey, A. L., El-Osta, A., Kingwell, B. A., Febbraio, M. A., Solari, F., Vidal, H. and Pirola L. (2011). Phosphoinositide 3-kinase as a novel functional target for the regulation of the insulin signaling pathway by SIRT1. Mol Cell Endocrinol. 335, 166–176.

32. Kulkarni, S. S. and Cantó, C. (2015). The molecular targets of resveratrol. Biochim Biophys Acta. 1852, 1114–1123.

33. Schrauwen, P. and Timmers, S. (2014). Can resveratrol help to maintain metabolic health? Proc Nutr Soc. 73, 271–277.

34. Choi, W. H., Son, H.J., Jang, Y.J., Ahn, J., Jung, C.H. and Ha, T. Y. (2017). Apigenin Ameliorates the Obesity-Induced Skeletal Muscle Atrophy by Attenuating Mitochondrial Dysfunction in the Muscle of Obese Mice. Mol Nutr Food Res. 61.

35. Skittone, L. K., Liu, X., Tseng, A. and Kim, H. T. (2008). Matrix metalloproteinase-2 expression and promoter/enhancer activity in skeletal muscle atrophy. J Orthop Res. 26, 357–363.

36. Schoser, B.G., Blottner, D. and Stuerenburg, H. J. (2002). Matrix metalloproteinases in inflammatory myopathies: enhanced immunoreactivity near atrophic myofibers. Acta Neurol Scand. 105, 309–313.

37. Carpéné, C., Gomez-Zorita, S., Gupta, R., Grès, S., Rancoule, C., Cadoudal, T., Mercader, J., Gomez, A., Bertrand, C. and Iffiu-Soltész, Z. (2014). Combination of low dose of the anti-adipogenic agents resveratrol and phenelzine in drinking water is not sufficient to prevent obesity in very-high-fat diet-fed mice. EUR J NUTR, 53,1625–1635.

38. Rossi, A. P., Rubele, S., Calugi, S., Caliari, C., Pedelini, F., Soave, F., Chignola, E., Vittoria Bazzani, P. Mazzali, G., Dalle Grave, R. et al. (2019) Weight Cycling as a Risk Factor for Low Muscle Mass and Strength in a Population of Males and Females with Obesity. Obesity. 27, 1068.

39. Wang, X., Zhao, D., Cui, Y., Lu, S., Gao, D. and Liu, J. (2019). Proinflammatory macrophages impair skeletal muscle differentiation in obesity through secretion of tumor necrosis factor-α via sustained activation of p38 mitogen-activated-activated protein kinase. Journal of Cellular Physiology. 234, 2566–2580.

40. Dupont-Versteegden, E. E., Knox, M., Gurley, C. M., Houlé, J. D. and Peterson C. A. (2002). Maintenance of muscle mass is not dependent on the calcineurin-NFAT pathway. Am J Physiol Cell Physiol. 282, C1387–95.

41. Adhikary, S., Choudhary, D., Tripathi, A. K., Karvande, A., Ahmad, N., Kothari, P and Trivedi, R. (2019). FGF-2 targets sclerostin in bone and myostatin in skeletal muscle to mitigate the deleterious effects of glucocorticoid on musculoskeletal degradation. Life Sci. 229, 261–276.

42. SanGiovanni, John, P. and Chew, E. Y. (2005). The role of omega-3 long-chain polyunsaturated fatty acids in health and disease of the retina. Progress in Retinal & Eye Research. 24, p87–138.

43. Iwata, Y., Ozaki, N., Hirata, H., Sugiura, Y., Horii, E., Nakao, E., Tatebe, M., Yazaki, N., Hattori, T. and Majima, M. et al. (2006). Fibroblast growth factor-2 enhances functional recovery of reinnervated muscle. Muscle Nerve. 34, 623–30.

44. Bennett, B. T., Mohamed, J. S. and Alway, Stephen E. (2013). Effects of Resveratrol on the Recovery of Muscle Mass Following Disuse in the Plantaris Muscle of Aged Rats. PLoS ONE. 8, 1–1.

45. Gliemann, L., Olesen, J., Biensø, R. S., Schmidt, J. F., Akerstrom, T., Nyberg, M., Lindqvist, A., Bangsbo, J. and Hellsten, Y. (2014). Resveratrol modulates the angiogenic response to exercise training in skeletal muscles of aged men. American Journal of Physiology: Heart & Circulatory Physiology. 307, H1111–H1119.

46. Bennett, B. T., Mohamed, J. S. and Always, S. E. (2013). Effects of resveratrol on the recovery of muscle mass following disuse in the plantaris muscle of aged rats. PLoS One. 8, e83518.

47. Le, N. H., Kim, C. S., Park, T., Park, J. H., Sung, M. K., Lee, D. G., Hong, S. M., Choe, S. Y., Goto, T. and Kawada, T. et al. (2014). Quercetin protects against obesity-induced skeletal muscle inflammation and atrophy. Mediators Inflamm. 2014, 834294.

